# Development of an EPR-based methodology to study protein-lipid interaction

**DOI:** 10.1101/2025.08.29.672991

**Authors:** Clara Piersson, Shikhar Prakash, Victoria Lublin, Melanie Rossotti, Baptiste Fischer, Madhur Srivastava, Yann Fichou

## Abstract

The interaction of protein with other biomolecules is central to all cellular processes. In particular, protein-lipid interactions play an essential role in regulating soluble and membrane protein function, structure, and dynamics. However, probing these interactions remains challenging due to the complexity and heterogeneity of membranes. Various methods have been developed to characterize protein-membrane interaction, each presenting advantages and limitations. This study presents a robust methodology based on continuous-wave Electron Paramagnetic Resonance (CW-EPR) spectroscopy to characterize protein–membrane interactions. We focused on the protein Tau, an intrinsically disordered protein associated with neurodegenerative diseases. We show that the interaction of labelled Tau with lipids gives rise to a very distinct lineshape, which can be used to quantify the fraction of bound protein. This allows to obtain the apparent binding mode and affinity through titration experiments. In addition, we show that a single measurement provides the absolute concentration of free and bound protein. We argue that this information, which is rarely obtained by other methods providing relative signals, is very useful for mechanistic studies. Furthermore, using mathematical modeling, we developed a minimal-data approach and demonstrated that a single EPR measurement can be used to derive binding constants. The approach is applied to the Tau-membrane interaction occurring in different conditions affecting the binding behavior. The presented methodology is expected to be applicable to other proteins.

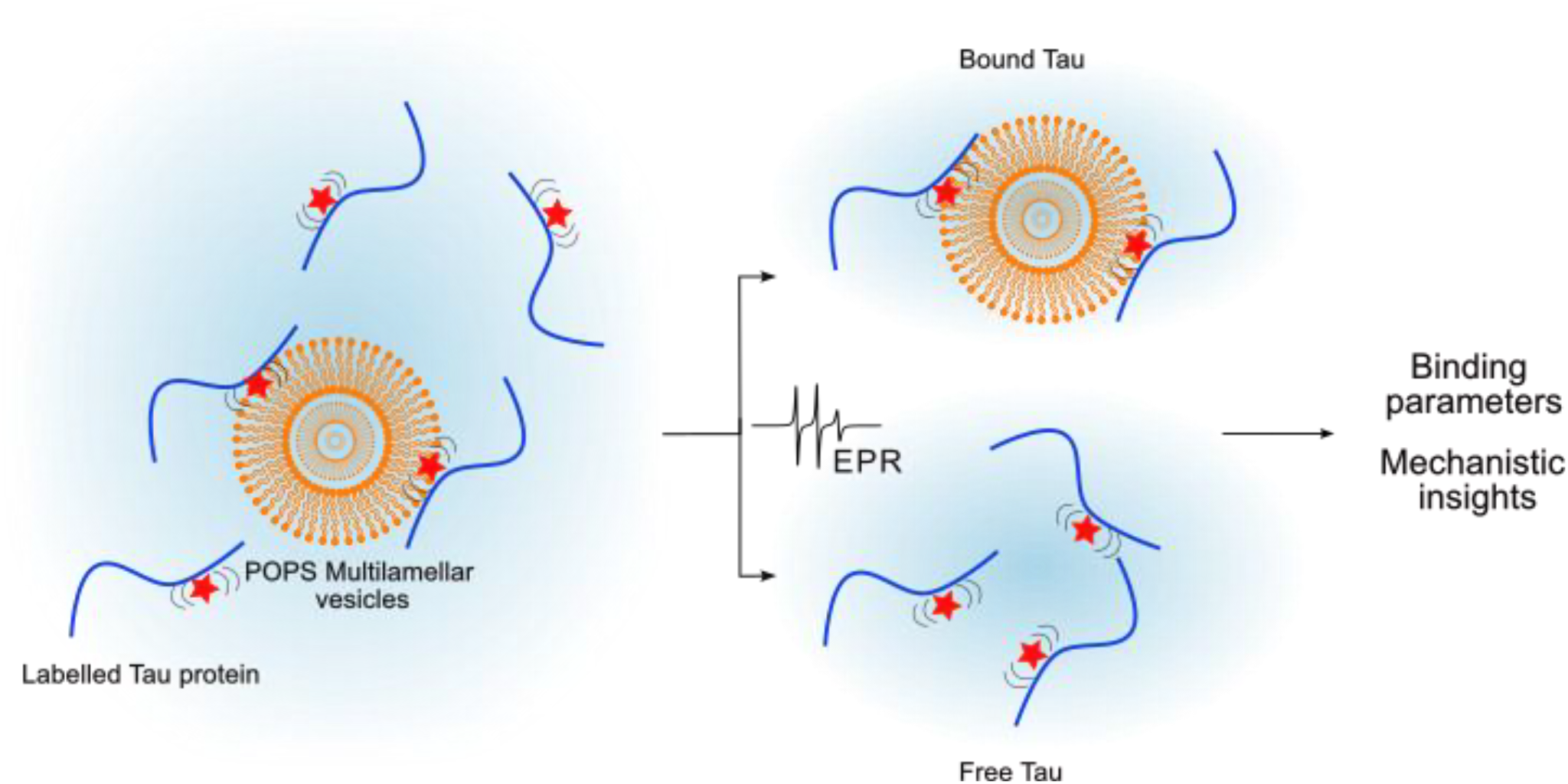

TOC Figure. Schematic representation of EPR-based analysis of Tau-membrane interactions using spin-labeled Tau and POPS vesicles

## A. Introduction

Binding affinity is a central aspect of biological function, as most physiological and pathological processes rely on interactions between biomolecules. Gaining insight into these interactions requires not only structural characterization, but also a quantitative understanding of the energetic forces driving complex formation. Over the years, a wide range of biophysical techniques have been developed to measure protein–ligand binding affinities, based on diverse principles such as calorimetry (isothermal titration, ITC), optics (fluorescence; Förster resonance energy transfer, FRET), diffusion (microscale thermophoresis, MST), interferometry (surface plasmon resonance, SPR; biolayer interferometry, BLI), and magnetism (nuclear magnetic resonance, NMR; electron paramagnetic resonance, EPR), among others. Each method offers specific advantages in terms of sensitivity, sample requirements, kinetic resolution, and compatibility with different systems and buffer conditions.

ITC measures heat changes upon ligand binding, providing direct and label-free access to thermodynamic parameters such as enthalpy, entropy and dissociation constants (K_D_) [1,2]. ITC is frequently applied in the characterization of protein-small molecule, protein-DNA/RNA, or protein-protein interactions.

Fluorescence spectroscopy techniques utilize fluorescent labeling to detect changes in emission intensity, polarization, or energy transfer efficiency upon ligand binding, allowing highly sensitive, real-time measurements even at low analyte concentrations [3]. In particular, FRET, which relies on the non-radiative transfer of energy between two fluorophores, can be used to quantify protein interaction [4]. MST enables sensitive, low-volume quantification of interactions in the pM to mM range by tracking the movement of fluorescently labeled molecules along a temperature gradient, which reflects binding-induced changes in size, charge, or solvation [5].

SPR and BLI are label-free techniques used to quantify binding affinities and kinetics in real time. SPR detects changes in refractive index near a sensor surface where the ligand is immobilized and analytes are flowed over via microfluidics [6]. Plasmon-waveguide resonance (PWR) is a notable variation of SPR spectroscopy that has proven powerful to characterize membrane-protein interaction [7]. BLI, in contrast, measures shift in light interference from an optical biosensor tip as analyte binds, without requiring flow systems. Both methods enable precise analysis of biomolecular interactions, with BLI offering greater simplicity and tolerance to complex samples [8].

Magnetic resonance techniques, including NMR and EPR, probe atomic environments and spin dynamics, offering structural and dynamic information simultaneously. NMR can be used to measure apparent affinity, typically by tracking chemical shift perturbation obtained with a ligand titration [9]. EPR is sensitive to unpaired electron found in radicals or metal centers and can provide insights into molecular dynamics, local environments, and intermolecular interactions [10]. By analyzing changes in the mobility of the spin-labeled molecule upon complex formation, EPR enables to characterize protein-protein interaction [11]. In particular, EPR has been used to study proteins-lipid interactions, describing for instance the location and the relative affinity of these interactions [12]. These studies have predominantly relied on spin-labeled lipids, enabling the precise characterization of the lipid environment and its perturbation upon protein binding [13].

In this study, we develop and implement an EPR-based methodology to quantitatively assess the interaction between the protein Tau and lipid membranes. Tau is an intrinsically disordered protein implicated in neurodegenerative diseases such as Alzheimer’s disease in which it forms highly ordered aggregates known as amyloid fibrils [4]. Beyond its canonical microtubule-binding function, Tau localizes near neuronal membranes [14] where its interaction with specific lipids might play a role in its pathological activity [15]. Tau is overall positively charged, whereas the POPS membranes studied here are negatively charged, creating favorable electrostatic interactions.

By monitoring changes in the mobility of a spin-labeled Tau variant upon lipid binding, we demonstrated that the binding modes and apparent binding affinities (*K*_*D*_) can be determined by titration. We further discuss how a single measurement, which provide the absolute quantity of bound protein, can be used to obtain apparent affinity. This article provides a robust framework for studying protein–membrane interactions with EPR in complex system.

## B. Methods

### 1. Tau expression and purification

Expression of the recombinant Tau 2N4R isoform harboring the P301L pathogenic mutation and a C291S substitution was performed in *E. coli* BL21(DE3) cells transformed with a pET28-based plasmid. A starter culture (10 mL) was grown overnight and used to inoculate 1 L of LB medium containing kanamycin (30 µg/mL). Cells were cultivated at 37 °C with orbital shaking (200 rpm) until reaching mid-log phase (OD_600_ = 0.6–0.8), at which point expression was triggered by the addition of 1 mM IPTG. After 3 hours of induction, cells were collected by centrifugation (5000 rpm, 20 min, 4 °C) and resuspended in a lysis buffer composed of 50 mM Tris-HCl (pH 7.4), 100 mM NaCl, and 0.1 mM EDTA, supplemented with 1mM PMSF, 5mM DTT, 20 µg/mL DNase, 10 mM MgCl_2_, and protease inhibitors. Lysis was carried out by lysozyme incubation (30 min, room temperature, agitation), followed by three freeze–thaw cycles using liquid nitrogen. After removal of cellular debris (9500 rpm, 10 min, 4 °C), the clarified lysate was subjected to a heat-treatment step (75 °C for 12 min), rapidly cooled on ice, and centrifuged again. The final supernatant was filtered and loaded onto a cation exchange column (UNO Sphere S, Bio-Rad), pre-equilibrated with 20 mM sodium phosphate buffer (pH 7.4) containing 0.1 mM EDTA and 100 mM NaCl. After washing, proteins were eluted using a linear gradient of NaCl up to 500 mM. Relevant fractions were pooled, concentrated, and further purified by gel filtration on a Superdex 200 Increase 16/600 pg column (Cytiva) equilibrated in 20 mM HEPES pH 7.4, 100 mM NaCl. The concentration of the purified Tau protein was determined spectrophotometrically at 274 nm using a molar extinction coefficient of 7.5 mM^−1^·cm^−1^. Samples were stored at –20 °C until use.

### 2. Liposome formulation into multilamellar vesicles (MLVs)

Multilamellar vesicles (MLVs) were prepared from 1-palmitoyl-2-oleoyl-sn-glycero-3-phospho-L-serine (POPS) alone or from mixtures of POPS and 1-palmitoyl-2-oleoyl-glycero-3-phosphocholine (POPC) with a 50/50 molar ratio. The lipids, initially dissolved in chloroform, were combined at the desired molar ratio and dried under a stream of nitrogen to form a thin lipid film. To ensure complete removal of residual chloroform, the film was then placed under vacuum in a desiccator overnight. The resulting dry lipid film was rehydrated in 20 mM HEPES buffer pH 7.4, leading to the formation of multilamellar vesicles.

### 3. Spin labeling of Tau

Tau mutant was labelled on cysteine residue (C322) using the nitroxide spin label MTSL (CAS 81213-52-7). Tau was first reduced with TCEP for at least 2 hours. TCEP was then removed using a PD-10 desalting column. A 10-fold molar excess of MTSL was added, and the mixture was incubated for a minimum of 2 hours under continuous stirring. Excess spin label was eliminated by two successive PD-10 columns. The concentration of spin-labelled Tau was determined using a standard curve generated with TEMPO (10 µM to 500 µM range), and signal integration (single and double integrals) was performed using SpinToolbox with a labeling efficiency of 60%.

### 4. Continuous-wave electron paramagnetic resonance (CW-EPR)

CW-EPR measurements were acquired using a Ciqtek EPR200M X-Band spectrophotometer operating at a microwave frequency of 9.8 GHz with modulation frequency of 100 kHz and a modulation amplitude of 1 G. The microwave power was set to 1 mW with an attenuation of 20 dB. The conversion time and time constant were set to 50 ms. The measurements were done at room temperature.

Samples were prepared by first mixing 50 µM labelled Tau with 0 to 25 mM of MLVs in 20 mM HEPES pH 7.4 buffer a few minutes prior to the measurement. Then, 6.4 µL were transferred in two 0.8 mm diameter quartz capillaries. 1D CW-EPR spectrum were acquired for 30 min. Spectra were fitted using SimLabel software [16], as detailed in the result section.

## C. Results

### 1. EPR spectral lineshape in the absence and presence of lipids

We performed site-specific spin labeling on tau 2N4R harboring the mutations P301L and C291S. The former is a disease-associated mutation whereas the later mutation removes one of the two native cysteines, enabling to only label the cysteine at position 322 with a nitroxide spin label MTSL. This mutant is referred to as Tau throughout the manuscript.

We measured the continuous-wave electron paramagnetic resonance (CW-EPR) spectrum of monomeric labelled Tau at a concentration of 50 µM (Fig 1B dark blue). The spectral lineshape exhibits narrow peaks, indicative of a highly mobile spin label, which is consistent with a nitroxide label attached to an intrinsically disordered protein [17]. We titrated Tau with increasing concentration of multilamellar vesicles (MLVs) composed of 1-palmitoyl-2-oleoyl-sn-glycero-3-phospho-L-serine (POPS) (Fig 1B blue to red). This titration shows that the spectral lineshape of monomeric Tau converges to a second well-defined lineshape at high lipid concentration (Fig 2B red), characteristic of restricted spin-label mobility. This change in lineshape reflects a direct interaction between Tau and POPS, as previously reported by other methods [18].

**Figure 1.**
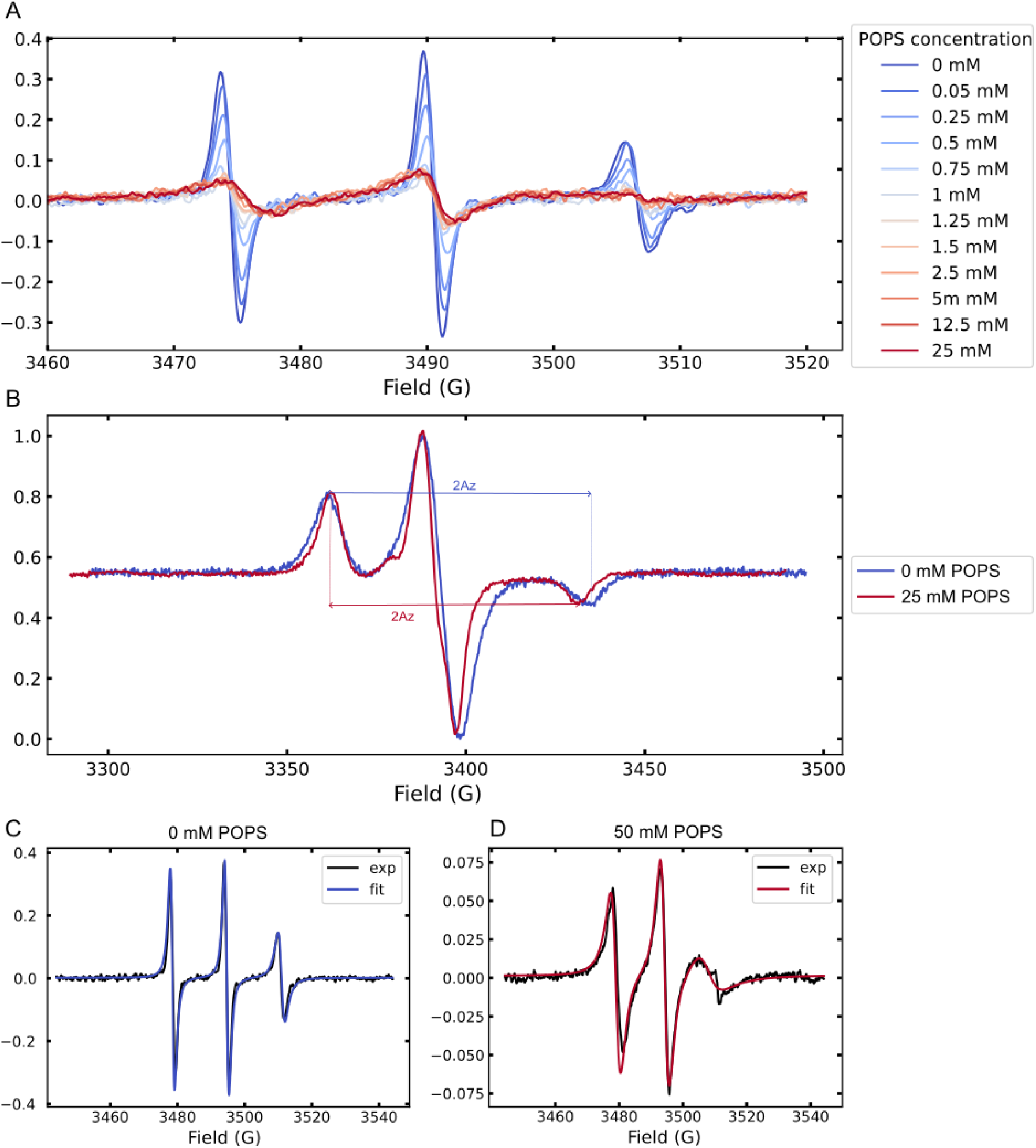
(A) CW-EPR spectra of labelled Tau at room temperature upon titration with increasing concentrations of POPS MLVs, showing progressive lineshape changes. (B) Frozen-state spectra (150 K) of labelled Tau in the absence (blue) and presence (red) of 25 mM POPS MLVs. (C) Spectrum of Tau monomer in a free environment at room temperature. (D) Spectrum of Tau monomer in a restricted environment at room temperature. [Tau] is 50 µM

**Figure 2.**
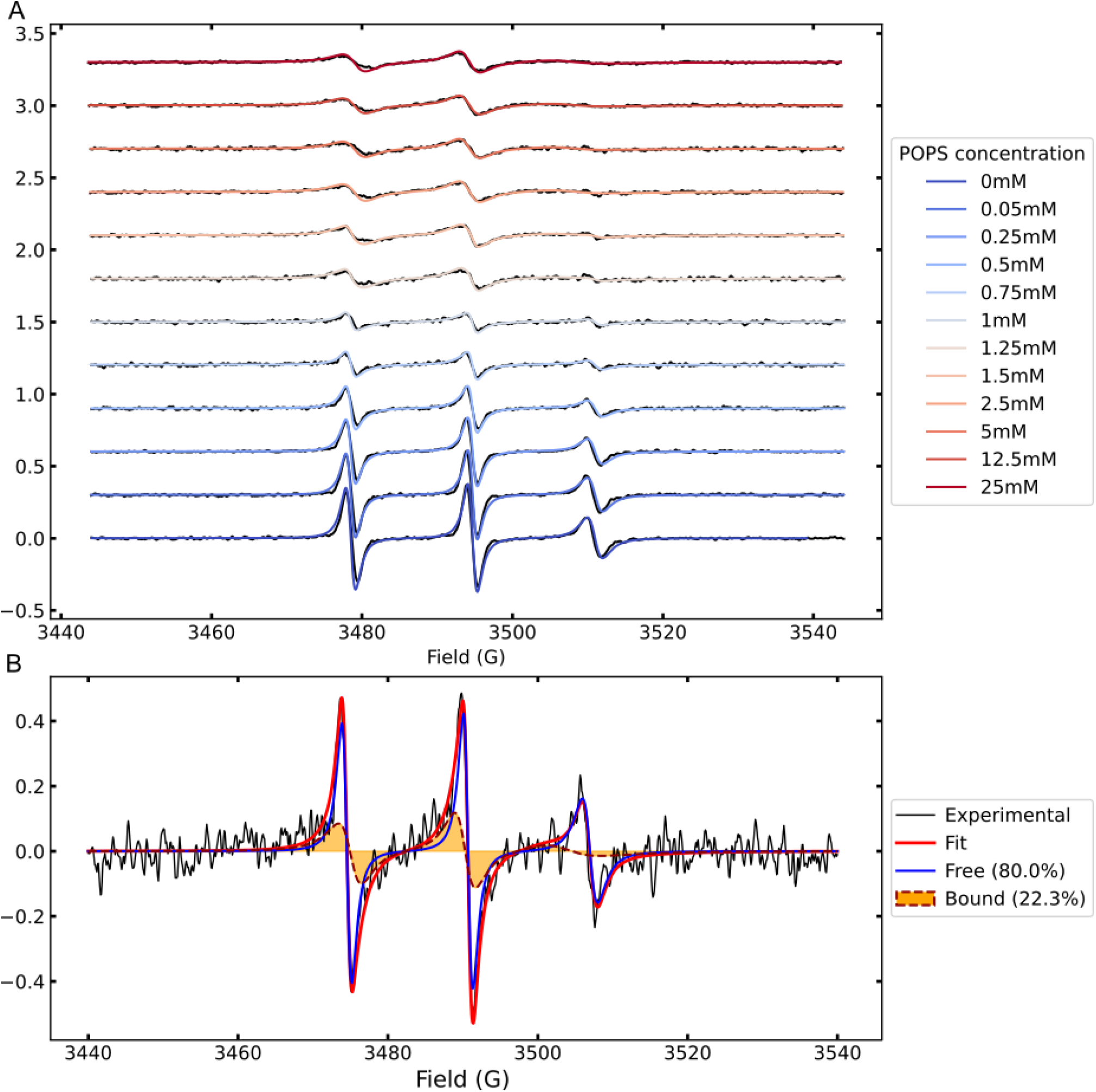
(A) Experimental data (black) and fit (blue to red) of labelled Tau at different POPS concentration. [Tau]=50 µM (B) Example of decomposition of experimental EPR spectrum (black) into the free component (blue) and bound component (orange fill), in the presence of 0.75 mM POPS, leading to satisfactory fit (red).

We fitted the experimental data corresponding to the first lineshape (Tau monomer) and the second lineshape (Tau in excess of POPS), using SimLabel software (Fig 1C & D) [19]. Following established guidelines, we first measured the *A*_*z*_ component from the spectra of frozen samples (Fig 2B) and determined *g*_x_ according to [19]:

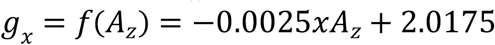

The g-tensor components (*g*_y_, *g*_z_) were fixed to 2.0061 and 2.0022 respectively following the guidelines [19]. The spectra were fitted by varying the correlation time, axial hyperfine coupling (*A*_*x*_=*A*_*y*_) and a Lorentzian line broadening (representing peak spacing). The fitted values for each condition are summarized in Table 1 and the corresponding simulation are shown in Figure 1 C,D.

**Table 1.** Fitting parameters obtained with Simlabel for Tau monomer and Tau bound to POPS. Fits are shown in figures 1C,D.

| *Fitting parameters* | *Tau Monomer* | *Tau bound to POPS* |
| --- | --- | --- |
| $g_x$ | 2.0083 | 2.0087 |
| $g_y$ | 2.0061 | 2.0061 |
| $g_z$ | 2.0022 | 2.0022 |
| $A_x = A_y$ (mT) | 0.60 | 0.60 |
| $A_z$ (mT) | 3.68 | 3.49 |
| *Correlation time (ns)* | 0.383 | 2.25 |
| *Lorentzian width (MHz)* | 0.12 | 0 |

### 2. Tau CW-EPR spectra with any lipid concentration can be decomposed into two components

Figure 2A shows the presence of two characteristic spectral lineshapes, with one progressively shifting toward the other upon addition of anionic lipids, POPS. These two components differ in their rotational correlation time (Table 1), which reflects the local dynamics of the nitroxide spin label. The first component corresponds to a fast-motion regime (Fig. 1C), while the second reflects slower dynamics (Fig. 1D). The fast component arises from unstructured, highly dynamic monomeric Tau in solution, whereas the slow component results from a direct interaction between spin label and a large object that restricts its motion, here POPS MLVs. Based on this, we define two distinct sates: monomeric Tau in solution, referred to as “free”, and Tau interacting with MLVs, referred to as “bound”. According to this definition, one expects that the titration of lipids should shift the populations of the free and bound states, which are in thermodynamic equilibrium.

Consistently with this prediction, we found out that all spectra can be accurately modeled using this two-component approach by simply changing the population of the free and bound components (Fig. 2). A representative example is shown in Figure 2B, where the individual contributions of the free (blue) and bound (orange) components are explicitly highlighted. From the above analysis, we extracted the relative proportions of bound and free Tau at each lipid concentration, allowing us to plot the fraction of bound protein as a function of POPS concentration (Fig. 3). This EPR-derived binding curve can then be fitted by conventional interaction model to obtain information on binding modes and constants, as described below.

**Figure 3.**
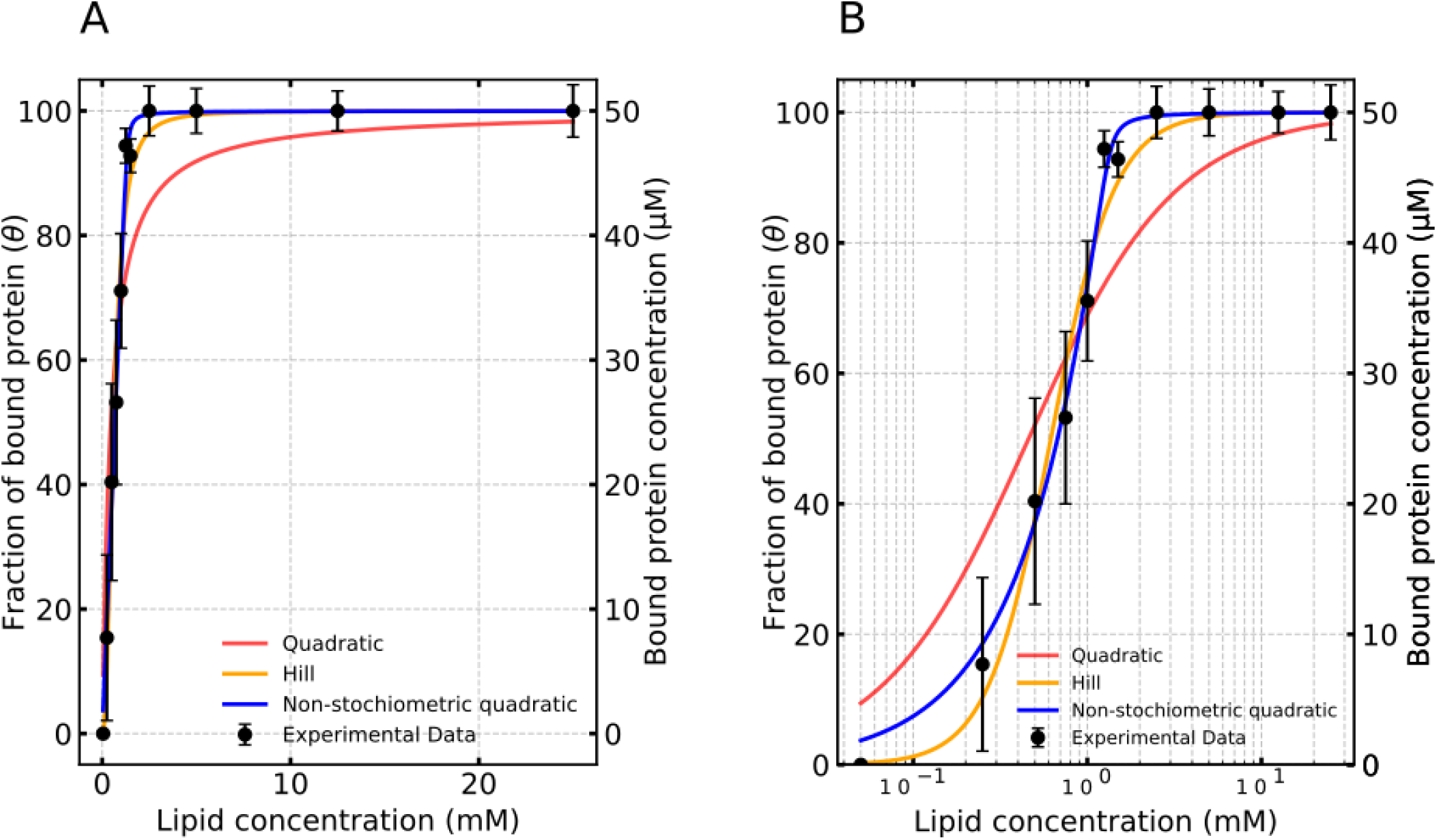
(A) Fit of experimental data on a linear lipid concentration scale, showing that binding saturates quickly at low millimolar concentrations. (B) Same dataset displayed on a logarithmic scale to better visualize the binding behavior at low lipid concentrations.

### 3. Fitting EPR-based binding curves

In order to elucidate the binding mechanism governing the interaction between Tau and POPS, the concentration of bound protein as a function of lipid concentration were fitted to different binding models: quadratic, non-stoichiometric quadratic, and cooperative binding models.

A quadratic model assumes a 1:1 binding stoichiometry between Tau and POPS, consistent with a simple, non-cooperative interaction [20]. It is described by the equation:

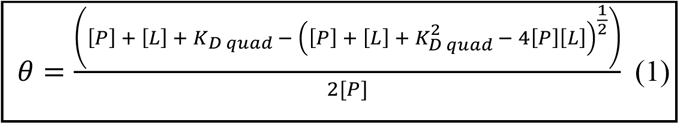

where [P] is the total protein concentration, [L] is the total lipid concentration and *K*_*D*_ is the apparent binding affinity constant.

The non-stoichiometric quadratic model indicates a non-linear behavior, where Tau binds to multiple POPS molecules (1:X stoichiometry, with X > 1). This model, described by the following equation, was previously used to describe protein interacting with multiple lipid molecules on a membrane [21].

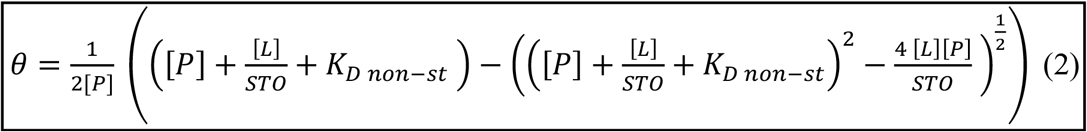

where *K*_*D non*−*st*_ refers to the apparent binding affinity and *STO* to the stoichiometry (*i.e*., number of interacting lipids per protein molecule).

Finally, cooperativity refers to a behavior in which the binding properties of each protein molecule are interdependent, *i.e*., initial binding events influence subsequent ones. It is often described by the simplified model of the Hill equation:

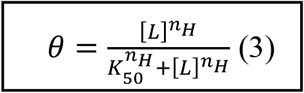

where *K*_50_ is the lipid concentration at half-maximal protein binding, rather than a true dissociation constant, due to the cooperative nature of the interaction. *K*_50_ is used in the Hill equation as an empirical affinity constant. *n*_*H*_ is a cooperativity parameter. The higher *n*_*H*_ is, the more cooperative is the system.

Figure 4 shows the fit of each model to the experimental data and the output fitted parameters are shown in Table 2. The data are best described by the non-stoichiometric quadratic model (blue) or Hill model (orange), indicating that our system tau-POPS is not optimally described by the simple quadratic model. The non-stoichiometric model, used previously for a very similar system [21], has the advantage of using a physical parameter directly interpretable, which is the number of lipid molecules interacting with one protein molecule (STO). However, as shown in Table 2, the variance of the apparent binding affinity is high (almost 100%), making this model intrinsically unprecise to determine the affinity. In contrast, the cooperative binding provides a more accurate affinity, although hill coefficient has no direct physical meaning.

**Table 2:** Fitting value achieved from different binding models. Uncertainties of the fitted parameters are extracted from the diagonal elements of the covariance matrix.

| Model | Apparant binding affinity<br>$K_D$ (mM) | Additional parameters |
| --- | --- | --- |
| Quadratic | $0.46 \pm 0.16$ | NA |
| Non – stoichiometric quadratic | $3.12 * 10^{-4} \pm 3.02 * 10^{-4}$ | $STO = 27.8 \pm 1.1$ |
| Cooperative | $0.64 \pm 0.04$ | $n_H = 2.30 \pm 0.04$ |

**Figure 4.**
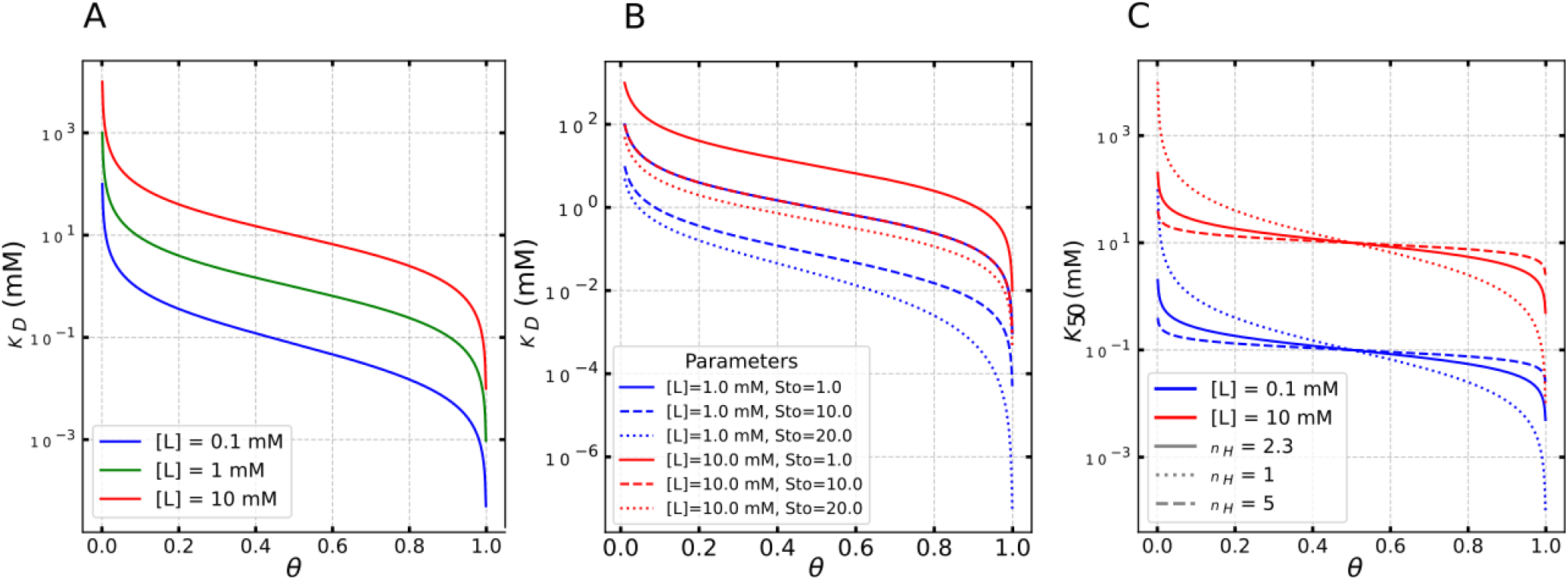
Relation between the affinity constants and the measured parameter *θ*, for (A) the quadratic (equation (4)), (B) the non-stoichiometric quadratic model (equation (5)) and (C) the Hill model (equation (6)). [*P*] is kept fixed at 50 µM and a few curves are plotted for different [*L*] and cooperativity parameters.

### 4. EPR provides absolute population quantification

The approach presented above exploits a typical binding curve, which can be built from any physical parameter that is proportional to the bound population (fluorescence, absorption, calories, chemical shift, etc… see introduction). The good fit of common interaction models (Figure 3) reinforces the finding that the two-component decomposition of the EPR spectrum, developed in section 2, can indeed probe the bound/free protein population.

However, the EPR-based population extracted here provides much richer information, which is the absolute concentration of bound protein. This contrasts with most methods used to investigate interactions that rely on relative readouts, which can only be interpreted through a full titration curve.

Indeed, since EPR is a quantitative method, every spectrum is decomposed into absolute concentrations of bound and free protein. This absolute quantification is particularly useful for mechanistic studies. For instance, directly assessing the absolute concentration of interacting protein is valuable for studies that aim to link a relevant mechanism (such as functional or pathological activity, aggregation, etc…) to the quantity of protein bound to a target. This capability is particularly advantageous when screening different environmental conditions (e.g., buffers, crowding) that can influence binding affinity. In such scenarios, EPR can directly assess the quantity of protein effectively bound in each condition, eliminating the need for a full affinity characterization through an entire titration curve.

The first implication of this absolute quantification is that binding affinity can be obtained from a minimal number of measurements, ideally one or two, according to equation (1)-(3). We specifically detail this approach below.

#### a. Measurement of affinity constants

For each model, *K*_*D*_ (or *K*_50_ in the case of Hill model) can be expressed analytically as a function of θ, the experimentally measured fraction of bound protein. These equations allow direct computation of the apparent binding constant from a single measurement.

For the quadratic binding model, equation (1) can be rearranged to isolate the square root

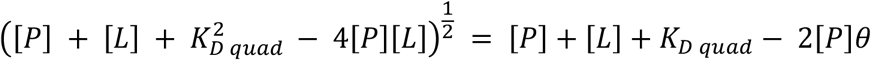

Squaring both sides and simplifying the equation:

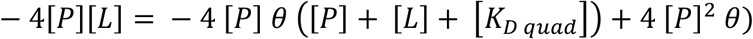

Finally, rearrange and solve to obtain *K*_*D quad*_ for quadratic model

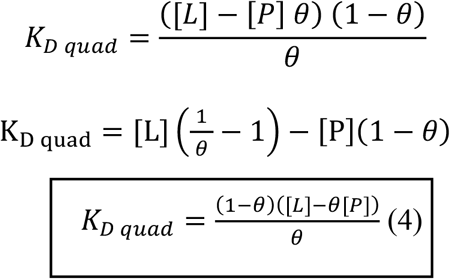

For the non-stoichiometric quadratic model, the *K*_*D non−st*_ can be extracted using by rearranging the equation. First isolating square root term from equation (2)

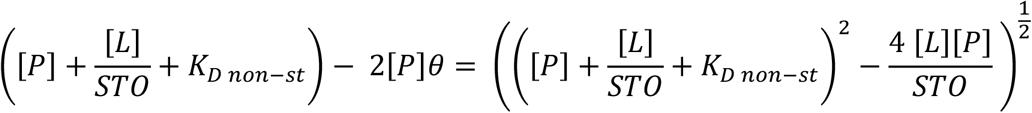

Squaring both sides and simplifying

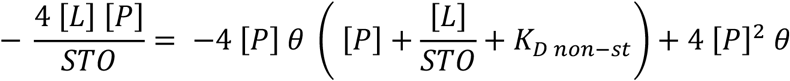

Rearranging and solving for *K*_*D*_

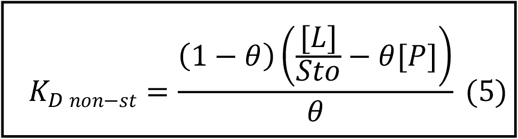

For the cooperative (Hill-type) model, the *K*_50_ is extracted by rearranging the equation (3)

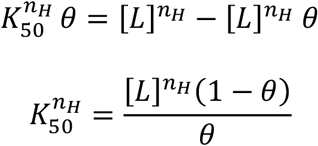

Therefore *K*_50_ for cooperative equation can be expressed as

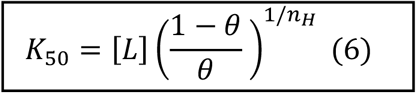

where *n*_*H*_ is the Hill coefficient. Note that in this model *K*_50_ is independent of [P] and is purely lipid-centric.

In these equations, [*L*] and [*P*], the concentrations of lipid and protein, respectively, are experimental input; θ, the fraction of bound protein, is measured by EPR ; and *K*_*D*_, *K*_50_, *n*_*H*_ and STO are unknown parameters characterizing the affinity of the protein and the lipid.

Figure 4 illustrates the relationship between the dissociation constant (*K_D_*) and the bound fraction (*θ*) for each model described by equations (4), (5) and (6). At the extremes of the binding curve, when *θ* approaches 0 (mostly unbound) or 1 (nearly saturated), the slopes of the curves are exceptionally steep. Consequently, even minor experimental uncertainties in θ in these regions can result in significant errors in the estimated *K*_*D*_.

In the Hill model, as shown in Figure 4C, curves corresponding to different values of *n*_*H*_ intersect at θ = 0.5. This implies that *K*_50_ can be determined independently of *n*_*H*_ when θ is measured near 0.5.

To further assess the reliability of single-point or minimal-point affinity measurements, we next derived analytical expressions for error propagation in each model. This analysis allowed us to identify the θ range where experimental uncertainty has the least impact on the calculated *K*_*D*_, thereby establishing a robust window for accurate affinity determination.

#### b. Analysis of error propagation

To evaluate the reliability of single-point *K*_*D*_ estimation across different models, we derived and analyzed the propagation of experimental error in the bound fraction θ to the calculated binding constants. Mathematically, this is expressed as the relationship between the relative error on 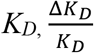, and the experimental uncertainty Δθ. For that, we defined a propagation factor A as 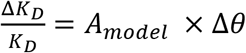.

For the quadratic model, the error propagation can be derived from the equation (4) by differentiating *K*_*D*_ with respect to *θ* :

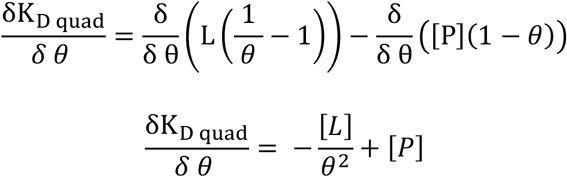

Dividing both sides by *K*_*d*_

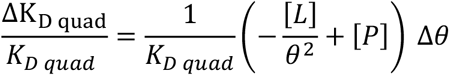

Substituting *K*_*D*_ from the equation (8) and solving for the final equation

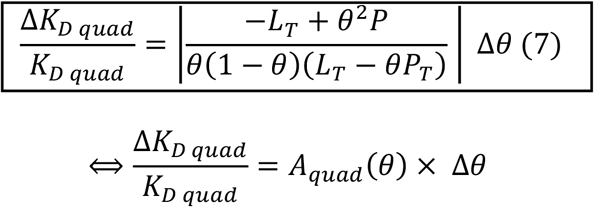

For the non-stoichiometric quadratic model, the expression of the error propagation factor A is obtained by differentiating *KD* with respect to *θ* from equation (5) to get 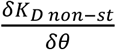.

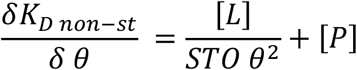

The error propagation formula for relative error is given by:

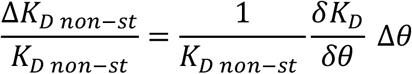

We get the final equation as :

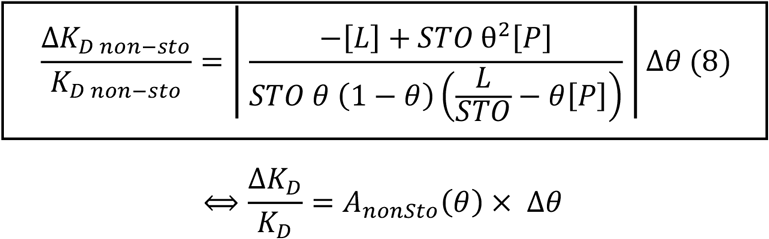

Finally, for the Hill model, the propagation can be calculated by first taking the log of equation (6) and then differentiating it with respect to theta:

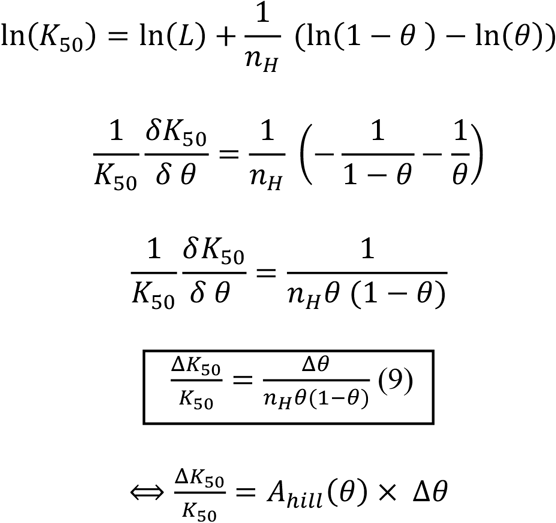

Our analysis of the error propagation factor, shown in figure 5, provides crucial insights into the precision of parameter determination. All models exhibit a U-shaped curve with errors increased drastically near θ = 0 and θ = 1, but minimized in the central region around θ = 0.5.

**Figure 5.**
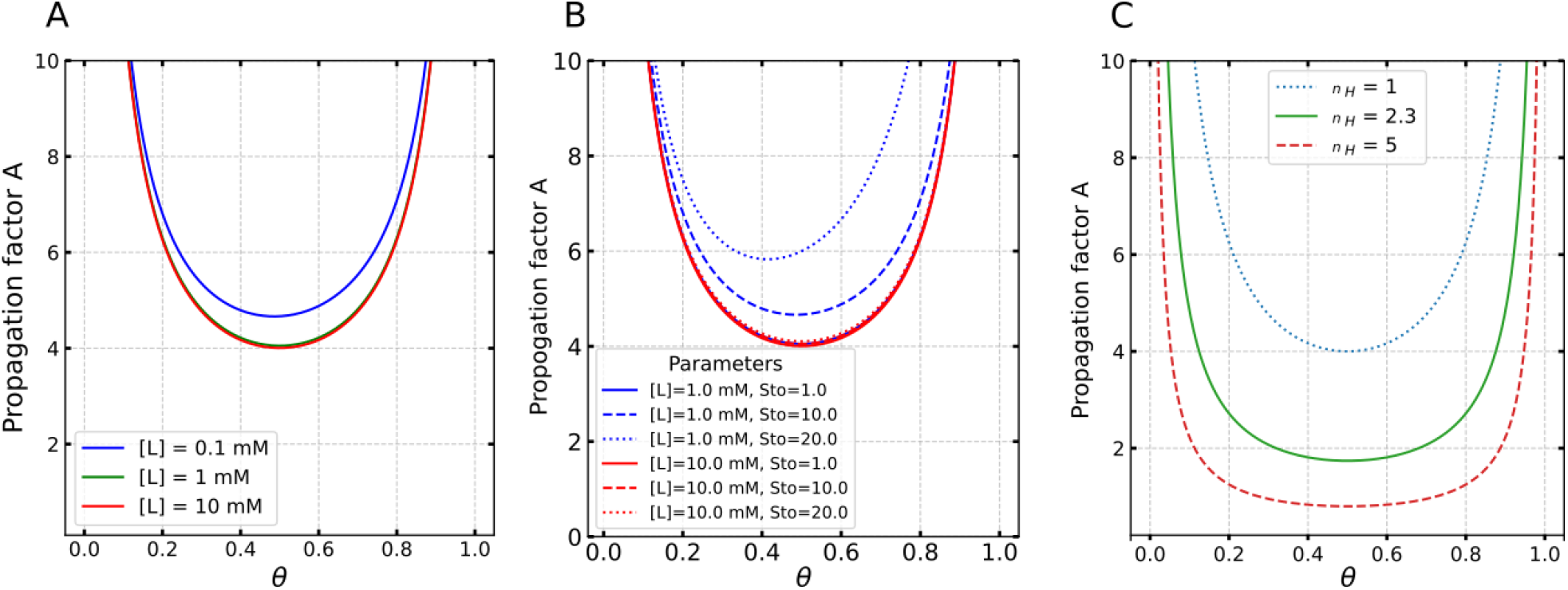
Error propagation factor A as a function of the bound fraction *θ* for three binding models defined by equations (7), (8) and (9). (A) Quadratic model: A is plotted for three total lipid concentrations ([L] = 0.1, 1, and 10 mM). (B) Non-stoichiometric model: A is calculated at two [L] values (2.7 mM and 278 mM) and three stoichiometries (STO = 10, 27.8, and 50). (C) Hill model: A is shown for Hill coefficients *n*_*H*_ = 1, 2.3, and 5. The minimum of each curve indicates the optimal θ region for minimal error in *K*_*D*_ estimation.

In the non-stoichiometric quadratic model (Figure 5B), the stoichiometry coefficient (STO) has a significant impact on error propagation. A higher STO is not favorable and increase the error propagation factor. In the Hill model (Figure 5C), the error propagation is inversely proportional to the Hill coefficient (n_H_), making the measurement of K50 more accurate in high cooperativity systems.

Altogether, these results support a general recommendation: to minimize uncertainty on affinity constants from minimal data, θ should ideally be measured within the 0.4–0.6 range, where error propagation is lowest.

#### c. Practical strategy for reliable *K*_*D*_ determination from minimal data

In practice, each EPR measurement at a given protein concentration [*P*] and lipid concentration [*L*] will provide an experimental value of *θ*. Determining the apparent dissociation constant *K*_*D*_ from a minimal number of data points requires the experimental approach to be adapted to the underlying binding model. Experimentally, the species containing the label, here *P*, is conveniently kept at a constant concentration, usually to compromise between sensitivity and protein consumption (typically tens of µ*M*). The concentration of lipid [*L*] is then the experimental parameter that can be changed to measure *θ* within the optimal range. If *θ* is too small (*i*.*e*., not enough bound protein), [*L*] can be increased. If *θ* is too large (*i*.*e*., too much bound protein), [*L*] can be decreased.

For the quadratic model, a single measurement of the bound fraction *θ* is sufficient as seen in equation (4). To ensure low error propagation, *θ* should be ideally measured between 0.4 and 0.6 (Figure 5A). Determining *K*_*D*_ in this region will be the most robust as it minimizes sensitivity to experimental uncertainty in *θ* measurements.

For the non-stoichiometric quadratic and Hill models, it is possible to determine the binding parameters (affinity and stoichiometry/cooperativity) from two measurements, as there are two unknown parameters (equation (5) and (6)). The model (non-stoichiometric quadratic or Hill) has to be assumed for the system under investigation.

The model can in principle be identified from this approach by measuring a third point. In that case the optimal conditions would be to measure one point at a lower concentration (around θ=0.25), one in the mid-range (around θ=0.5), and one at a higher concentration (around θ=0.75). Yet, this minimal data approach is not ideal when the binding model is unknown.

If the binding mode is unknown, the most pragmatic strategy is to perform a single measurement around *θ* = 0.5. First, this leverages the fact that, across all models, error propagation is minimized at this value (Figure 5). Moreover, figure 4 shows that around θ = 0.5, the *K*_*D*_ determination is independent of the model between Hill and quadratic models (recalling that *n*_*H*_ = 1 in the hill model is equivalent to the quadratic model). In that case, the minimal-data approach would be used for qualitative or relative assessment of the affinity. Recommendations are summarized in table 3.

**Table 3.** Recommended strategy for minimal-point *K*_*D*_ estimation based on the binding model

| Binding model | Minimal number of $\theta$ values | Recommended $\theta$ values | Parameters determined | Practical notes and limitations |
| --- | --- | --- | --- | --- |
| Quadratic (1:1) | 1 | 0.4–0.6 (ideally 0.5) | $K_D$ | Most robust estimation. Single-point measurement in this range minimizes error propagation (see Fig. 6). |
| Non-stoichiometric quadratic | 2 (for parameters), 3 (for model identification) | Within 0.2–0.8 (ideally 0.25, 0.5 and 0.75) | $K_D, STO$ | Two points allow solving for both parameters analytically. Requires a known model. Error propagation increases with higher $STO$. |
| Hill (cooperative) | 2 (for parameters), 3 (for model identification) | Within 0.2–0.8 (ideally 0.25, 0.5 and 0.75) | $K_{50}, n_H$ | Two points suffice to estimate parameters; three are needed to distinguish cooperative vs. non-cooperative behavior. |
| Unknown model | 1 | $\approx 0.5$ | Approximate $K_D$, | Best guess strategy. Minimal error at $\theta \approx 0.5$ across models (Fig. 6). However, result is only valid if underlying model is Hill or quadratic. High risk of model misinterpretation. |

### 5. Application physiological conditions

Building on the minimal-data approach to estimate apparent dissociation constants, we applied this methodology to two different data sets of the Tau/lipid system. In addition to the conditions used in Figure 4 (membrane POPS), we performed measurements with a modified membrane composition (membrane POPS:POPC 50:50). We measured the population of bound Tau θ at three different lipid concentrations. The data are summarized in Table 4.

**Table 4.** Population θ of bound Tau measured at three different lipid concentration for two different membrane compositions (100 % POPS and 50/50 % POPS/POPC) and binding parameters obtained from different approaches. * denotes the data used to calculate K50 from single data point. To determine K50 from single data point, *n*_*H*_ was fixed to the value obtained from all data. Data for POPS membrane are selected from the dataset presented in figure 4. Binding parameters from complete titration are obtained from the fit shown in figure 4.

| | Experimental data :<br>($\theta$, [L] (mM)) | $K_{50}$ from<br>complete<br>titration | $n_H$ from<br>complete<br>titration | $K_{50}$ from<br>three points | $n_H$ from<br>three points | $K_{50}$ from a<br>single data<br>point |
| --- | --- | --- | --- | --- | --- | --- |
| POPS | (0.40±0.16, 0.5) | 0.64 ±0.04 | 2.30 ±0.04 | 0.65±0.04 | 1.81±0.04 | 0.72 ± 0.16 |
|  | (0.53±0.13, 0.75)* |  |  |  |  |  |
|  | (0.71±0.09, 1) |  |  |  |  |  |
| POPS:POPC | (0.41, 16, 0.5) | / | / | 0.67±0.01 | 1.26±0.03 | 0.78 ± 0.15 |
|  | (0.58, 0.11, 0.875)* |  |  |  |  |  |
|  | (0.69, 0.6, 1.25) |  |  |  |  |  |

It is worth noting that, although measuring three distinct points around θ = 0.25, 0.5 and 0.75 is theoretically optimal to obtain reliable binding parameters, it is experimentally challenging to precisely obtain these θ values.

We applied the cooperative binding model that gave the best fit in Figure 4. We fit three data points around 0.25,0.5 and 0.75 to equation (6) and extract the corresponding K_50_ and *n*_*H*_. We obtained K_50_ = 0.65 ± 0.04 mM when three points are used, as compared to 0.64 ± 0.04 mM found for the complete dataset (table 4). Both values are identical within error, indicating that the three-point approach provides a robust and reliable estimate of K_50_. A more pronounced difference is observed in the Hill coefficient (*n*_*H*_), which reflects binding cooperativity. *n*_*H*_ of 1,81±0.04 was found with three points, as compared to 2.30 ± 0.04 obtained using all data (table 4). Although both values remain close to 2, this difference shows that the minimal-data approach is less robust to evaluate cooperativity.

Then, we estimated the apparent dissociation constant using a single data point around θ ≈ 0.5. We fixed *n*_*H*_ to the one obtained with all data and we extracted K_50_ and its uncertainty from equation (6) and (9), respectively. K_50_ was found at 0.71 ± 0.16 mM, exhibiting a remarkable similarity with K_50_ obtained from the full titration (0.64 ±0.04). The uncertainty is larger (22% for single point determination versus 6% for full titration), but remains within an acceptable range.

We applied the same approach for the membrane composed of 50/50 % POPS/POPC. From three data points, we could also obtain both K_50_ and *n*_*H*_, taking the values of 0.67 ± 0.01 mM and 1.26 ± 0.03, respectively (table 4). While the affinity is similar than for POPS membrane, *n*_*H*_ was found significantly smaller, indicating less cooperativity when the membrane charges are reduced. Finally, with a single data point, fixing *n*_*H*_ to the value obtained with more data, we found a K_50_ of 0.78 ± 0.15 mM. Similarly to the POPS membrane model, K_50_ obtained from a single point is similar to the one obtain from more data (0.78 ± 0.15 mM, compared to 0.67 ± 0.01 mM) and only the uncertainty increases (19% uncertainty with a single data point). Altogether, these results demonstrate that we can obtain reliable estimate of binding parameters with minimal experimental points: 3 data points allows to measure both K_50_ and *n*_*H*_ while a single point provides K_50_ with about 20% uncertainty if *n*_*H*_ is known.

## D. Conclusion

In this study, we developed and validated a CW-EPR–based methodology to quantify the apparent binding affinity between an intrinsically disordered protein and lipid membranes. By performing a spectral decomposition of spin-labeled Tau and carefully modeling the free and bound populations, we demonstrated that we can reliably extract the absolute concentration of bound and free protein.

From the population of bound protein, we showed that we can fit different binding models, quadratic, non-stoichiometric, and cooperative. In our conditions, we demonstrated that the Hill model provides the most reliable description of the Tau–lipid interaction, offering well-defined parameters and capturing potential multivalent or clustering effects at the membrane surface.

Furthermore, we established a framework for estimating apparent binding affinity from a single or limited number of experimental data points. This is possible by leveraging the fact that EPR-extracted population provides the absolute concentration of bound protein. We provide practical consideration to minimize uncertainty when using the minimal-data approach.

Altogether, this work introduces a versatile and accessible EPR-based approach for probing protein–membrane interactions. It paves the way for future studies investigating the molecular determinants of membrane recruitment, particularly in the context of intrinsically disordered proteins and their role in pathological aggregation processes. We also expect this approach to be applicable to other system (interaction between any protein and membrane, interaction between proteins), provided that the bound state leads to a change in EPR lineshape.

## E. Acknowledgements

The authors thank the European Research Council (Grant 101040138), the Fondation Vaincre Alzheimer and the Federation of European Biochemical societies for their financial support.

